# A High-Quality Acetylation Dataset Reveals Modest Data Requirements for Transfer Learning to Identify Little Studied Post-Translational Modifications

**DOI:** 10.64898/2026.06.25.733353

**Authors:** Yannick Luplow, Shengbo Wang, Andrew R. Jones, Juan Antonio Vizcaíno, Christoph N. Schlaffner, Bernhard Y. Renard

## Abstract

Dysregulation of post-translational modifications (PTMs) is associated with severe pathologies, including cancers and Alzheimer’s disease. Despite their biological importance, identifying modified peptides remains challenging due to the immense combinatorial search space. While searches benefit from prior knowledge of a peptide’s modification status, the data scarcity for most PTMs hinders the development of accurate deep learning classifiers like AHLF (ad hoc learning of peptide fragmentation). Here, we overcome this data bottleneck for acetylation and ubiquitination. We harmonised a dataset with about 500,000 high quality acetylated peptide-spectrum matches (PSMs) from nine publicly available acetylation-enriched datasets. We fine-tuned AHLF with the acetylation and a 2-million spectra strong ubiquitination dataset separately and assessed the minimum data requirement for training by iteratively downsampling. Training separate models on SILAC and label-free subsets also assessed the impact of data diversity. The resulting acetylation and ubiquitination models achieve an AUC of 0.87 and 0.90 respectively. Beyond 28,500 acetylated spectra, corresponding to roughly 0.3% of the original model’s training data, additional data just provides minor performance gains. Finally, we show that data diversity is beneficial for generalizability, while models trained on homogeneous data sources tend to overfit to their respective data type. All code, and model weights are available at https://gitlab.com/dacs-hpi/ahlf-ptmai.

## Introduction

Proteins are the primary executors of cellular function. Rather than acting as static end-products of gene expression, their activity, localization, stability, and molecular interactions are continuously modulated through post-translational modifications (PTMs). Dysregulation of PTM networks has been implicated in a wide range of pathological conditions, including cancer, diabetes, and Alzheimer’s disease [1–3]. Using proteomic mass spectrometry, over 650 distinct PTM types have been described to date [4]. They span a broad chemical spectrum, from small groups such as phosphorylation, to the covalent attachment of entire proteins [5]. Thereby, they vastly expand the functional complexity of the proteome. Recent advances in mass spectrometry-based proteomics have dramatically expanded our ability to characterize the PTM landscape of the proteome [6]. However, the computational identification of PTMs from mass spectrometry data remains a fundamental challenge. Conventional database search strategies are constrained to a limited set of predefined modifications and specific sequence windows, as the simultaneous consideration of all possible PTM combinations leads to a combinatorial explosion of the search space that renders exhaustive search computationally intractable [7, 8]. Also in recent years, the rapid advancement of deep learning has had a profound impact on proteomics, particularly in the area of de novo peptide sequencing [9–11]. Unlike database search approaches, de novo models identify peptide sequences directly from spectra without relying on a predefined sequence database [12]. Beyond de novo sequencing, deep learning has also been applied to spectrum classification, as demonstrated by AHLF (ad hoc learning of peptide fragmentation) [13]. AHLF is a binary classifier trained to distinguish phosphorylated from unphosphorylated spectra, illustrating the ability to learn spectrum representations and enable the stratified identification of PTMs. While deep learning models have made substantial progress in peptide identification from spectra, the training of such models requires massive amounts of data and the performance highly depends on the quality of the training data [14]. Commonly available datasets, which were used to train the aforementioned models, differ substantially in their composition. On the one hand, MassIVE-KB v1 is a diverse collection of 30 million publicly available real-world spectra for 2.1 million peptides and peptidoforms [15]. On the other hand, the ProteomeTools subsets MULTI-PTM [16] and 21-PTM [17] of synthetic peptides with 14 and 21 distinct PTMs consist of a total of 19.6 million and 1 million spectra [18], respectively. However, these well-defined datasets represent a static snapshot and do not scale with the continuously growing public proteomics repositories [19], leaving formerly underrepresented or unreported PTMs beyond phosphorylation largely untouched. This is best illustrated by AHLF, whose training relied on a reference phosphoproteome comprising roughly 10.5 million phosphorylated and 8.7 million unphosphorylated spectra [20]. Such a dataset harmonisation was only possible as phosphorylation is the most-studied PTM [21]. The available data for other biologically relevant PTMs such as acetylation and ubiquitination [22–24] remains rather scarce, limiting the extension of deep learning models to different modification types [11]. To address this scarcity, we created a high-quality harmonised dataset for human acetylation from nine publicly available acetyl-enriched datasets including 840,000 modified spectra. We further reused a 2.4 million spectra strong similarly harmonised human ubiquitination dataset [25] to apply transfer-learning to the phosphorylation pre-trained AHLF model, thereby, bypassing the requirement for large datasets to fully train models from scratch. To further evaluate the data requirements of the fine-tuning approach, the effect of training set size was systematically assessed through iterative downsampling. Additionally, the impact of data cleanliness on model performance was evaluated by comparing models trained on SILAC-labeled versus label-free spectra.

## Methods

### Dataset curation and preparation

#### Selection of datasets

To harmonise and generate a deep-learning-ready acetylation dataset, we selected datasets from the PRIDE repository for re-analysis based on the following criteria: (i) human-derived samples enriched for acetylation; (ii) data generated using Thermo Fisher Scientific instruments; and (iii) availability of metadata, either through the original publication or by direct communication with the authors. After the preliminary curation, 25 acetylated datasets were identified. Of these nine sets met the selection criteria and were retained. Among these nine acetylation-enriched datasets, four were chemically or other induced: one involved aspirin-acetylated proteins, another was chemically induced by NAS-d3, a third used acetyl-phosphate, which can actively acetylate proteins by itself (with SILAC heavy samples as induced and SILAC light samples as non-induced), and the fourth involved C- and D-acetic anhydride as an acetylating agent. A summary of the datasets, including PRIDE accession numbers, biological samples, and key characteristics, is provided in Table 1 for Acetylation datasets.

**Table 1:** List of Acetylation-enriched proteomics datasets and their main characteristics.

| Datasets | #raw files | Instruments | Label | Chemical/<br>Biological conditions | Biological samples | PubmedID |
| --- | --- | --- | --- | --- | --- | --- |
| PXD003530 | 24 | Q Exactive | label-free | Aspirin induced | HeLa Cells | 27913581 [26] |
| PXD005252 | 52 | Q Exactive | SILAC | Protein-induced | Kasumi-1, 293FT cells | 29804834 [27] |
| PXD005903 | 87 | Q Exactive | label-free | NAS-d3 induced | SiHa and Calo | 28893905 [28] |
| PXD009839 | 32 | Q Exactive | label-free | HCMV AD169 infection | lung fibroblasts | 30470744 [29] |
| PXD009994 | 123 | Q Exactive | SILAC | SILAC heavy chemical induced | HeLa Cells | 30837475 [30] |
| PXD023084 | 61 | Q Exactive HF-X | label-free | No induction | muscle biopsies | 35638262 [31] |
| PXD023218 | 32 | Q Exactive HF | dSILAC/TMT | No induction | HeLa Cells | 35013197 [32] |
| PXD041171 | 12 | Q Exactive HF-X | label-free | HCMV AD169 or TB40/E + AGK2 SIRT2 inhibitor | lung fibroblasts | 37916830 [33] |
| PXD014453 | 18 | Orbitrap Fusion Lumos | label-free | C- and D-acetic anhydride induced | MCF7, EK293 cells | 32290654 [34] |

#### Proteomics raw data processing

Datasets were analysed separately, raw files from each dataset were converted to mzML format using ThermoRawFileParser (version 1.3.4) [35] and analyzed independently. An initial subset of raw files from each dataset was processed using Fragpipe (https://fragpipe.nesvilab.org/) with an open search to identify modifications. Modifications detected in more than 1% of peptide-spectrum matches (PSMs) were retained for downstream analysis. Peptide and protein identification, including PTMs, was performed using the Comet search engine (version 2024) [36] on a Linux-based high-performance computing cluster. Default parameters were applied, with the following exceptions: missed cleavages were set to 4, and variable modifications were based on Fragpipe results, with oxidation of methionine and N-terminal protein acetylation (excluding N-terminal peptide acetylation) included for all the datasets. The search database consisted of the UniProt human reference proteome (one protein per gene, downloaded on April 4, 2024) and a contaminants database (https://www.thegpm.org/crap/, obtained April 2024). Decoys were generated using the reverse decoy method to the FASTA file via FragPipe. Statistical validation of PSMs and distinct peptide sequences was conducted using PeptideProphet [37] and iProphet [38] from the Trans-Proteomic Pipeline (TPP version 7.1.0) [39]. High-confidence PSM matches were obtained, and PTM site localization was computed using PTMProphet (TPP) [40], generating a unified mzidentML format file.

#### Post-processing

The search result files from the TPP were processed using a custom Python script (mzidFLR; https://github.com/PGB-LIV/mzidFLR), as previously described [41] and also applied in prior PTMeXchange projects [42]. First, a global false discovery rate (FDR) was calculated at the PSM level at a 1% threshold to retain high-confidence matches. The results were converted to a site-based format, with PTM localization scores assigned to each acetylation site. Contaminants, decoy hits, and non-acetylated PSMs were excluded, retaining only acetylated sites for downstream analysis. Second, to estimate the probability of correct PTM localization, the PTM localization probability (from PTMProphet) was multiplied by the PSM probability (from PeptideProphet). Redundancy in PSMs site-based evidence was addressed by collapsing the data to the peptidoform level using a binomial adjustment [43]: the probability of a site being acetylated was calculated by comparing the number of acetylation events to the total evidence for that site across all PSMs. False localization rates (FLRs) were estimated by introducing decoy alanine residues, yielding a 1% FDR PSM file and a PTM site-specific FLR file for quality control (QC) analysis for each individual dataset. To combine results across datasets, a meta-analysis approach was employed to control FLR inflation, referred to as PTM stats in meta-analyses. This approach addressed the accumulation of FDR due to the identification of the same sites across multiple datasets and the non-comparability of PTM probabilities across studies. PTM sites were classified into Gold, Silver, and Bronze categories. Classification thresholds were set as follows: Gold (threshold *≥* 2 datasets with <1% global FLR) and Silver (threshold = 1). A similarly harmonised dataset for ubiquitination was downloaded from PRIDE (PXD:PXD068989) [25] comprising eleven publicly available ubiquitination-enriched datasets.

#### Data Preparation to apply Transfer Learning

To further prepare the data for transfer-learning all raw files were converted to MGF format using ThermoRaw-FileParserGUI (v1.7.4), selecting only MS2-level spectra. We assigned each FLR category to the PSMs by the modified peptide sequence and applied the following filtering steps: (i) removed all duplicated scan IDs; (ii) removed all PSMs with PTMs on the C-terminus (since acetylation and ubiquitination events block tryptic digestion, these are likely false positive detections); (iii) removed all PSMs which mapped to decoy sequences; (iv) removed all PSMs which mapped to multiple proteins; (v) removed all PSMs with an FDR >1%; and (vi) removed all modified PSMs with an FLR category of Bronze. The remaining PSMs from each dataset were then concatenated into a single csv file containing information about the Protein ID, Modification status, FLR category, Universal Spectrum Identifier (USI), start and stop position of the peptidoform within the protein sequence. The corresponding spectra for all remaining PSMs were subsequently extracted from the MGF files.

### Split of the data into a Training, Validation and Test set

In order to apply transfer learning to the phosphorylation-pretrained AHLF model, we split our data into training, validation, and test subsets. To ensure that overlapping sequences were not distributed across different subsets we mapped peptide sequences back to their protein sequences and grouped overlapping peptides together into contigs, as illustrated in Figure 1. Thereby 11% of the acetylation and 12% of the ubiquitination peptides map to multiple proteins, were disregarded for contig generation and removed from further analysis. Contigs were then used as the unit of splitting, ensuring that overlapping peptides were sampled into the same subset. For the acetylation dataset we performed a rough 85/5/10 and for the ubiquitination dataset a rough 89/1/10 split. We further applied two different sampling methods. In the first approach, all modified peptides determined the total dataset size, the unmodified peptides were then randomly sampled from the same data split as the modified peptides; and if insufficient to balance the dataset contigs were added where no modified peptide is present. Subsequently the unmodified peptides were sampled to an equal amount of modified peptides within the corresponding set (split_unmodsampled). In the second approach, equal numbers of modified and unmodified spectra were sampled from the same contig (split_fullsampled). For both sampling techniques, only PSMs with a Gold label were included in the test set, ensuring the highest possible data quality for evaluation.

**Figure 1:**
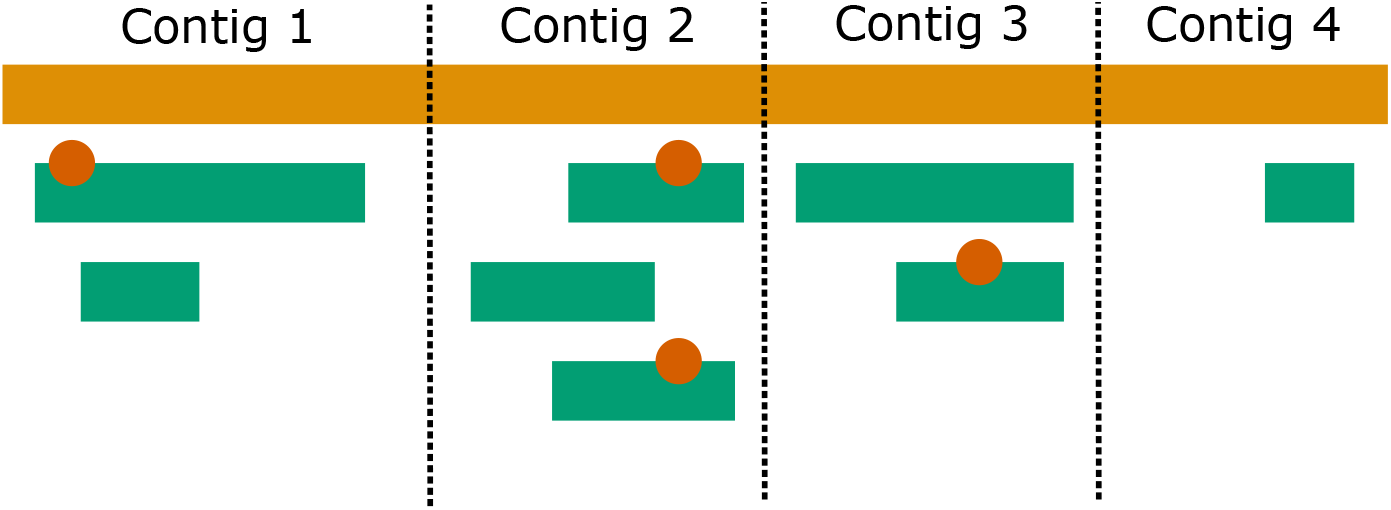
Sampling and splitting strategy to prevent data leakage. Peptide sequences (green bars) are mapped to the originating protein sequence (orange bar). Red circles indicate PTMs on the peptide sequences. Contigs are split into training, validation and testing datasets as contig 2, contig 1, and contig 3, respectively. The illustration further highlights the two sampling methods. In the first method, all modified peptides from a set are used, for example both modified peptides from contig 2 and the additional contig 4 is used to balance the training set between modified and unmodified peptides. In the second method, only one of the modified sequences is selected to ensure that the training set is balanced.

### Model training and assessment

We applied transfer learning by fine-tuning the AHLF alpha model, which was originally trained on phosphorylation, separately for acetylation and ubiquitination. To do so we froze all layers, except the last three dense layers and continued training for at most 40 epochs. After each epoch a validation step was performed to monitor the early stopping criterion based on the validation loss. A batch size of 128 and 256 was used for acetylation and ubiquitination, respectively. The initial learning rate was set to 1e-3, and the dropout rate was set to 3e-1 for both use cases. To ensure that model performance was not dependent on a particular data split, fine-tuning was performed for each of the five fullsampled data splits, and once, as it utilizes the maximum available training data, for the unmodsampled data split described above. All training and testing was performed on a single Tesla V100-SXM2-32GB GPU. The model performance was evaluated using scikit-learn (version 1.6.1) with balanced accuracy (Bacc), ROC-AUC, and F1-score as metrics.

### Assessing the data requirement

To assess the required amount of data for effective fine-tuning, we randomly downsampled the data for training and validation iteratively by half, resulting in ten dataset sizes ranging from the full dataset down to 1/512th of the complete data split. The training process was performed as described above for each of the downsampled sets and the resulting models were evaluated on the unchanged full test set. For each metric (Bacc, ROC-AUC, F1) a power law function (Equation 1) with *a ∈* [0.5, 1], *b ∈* [0, 1], *c ∈* [0.01, 2] was fitted to the performance values across dataset sizes.

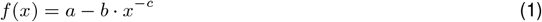

To account for the higher reliability of larger dataset sizes, a weighted least squares fitting was applied using weights, such that larger dataset sizes contributed more strongly to the optimization. To estimate 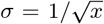 uncertainty, confidence intervals were computed using 300 bootstrap iterations with replacement. Lower and upper confidence bounds were set as 2.5th and 97.5th percentiles. The elbow point for each fitted curve was identified using the python package kneed (version: 0.8.6) [44]. The average across the three metrics was calculated to report a final elbow point. Analysis for acetylation and ubiquitination was performed separately.

### Assessing the data diversity

Since the curated datasets originated from studies using either SILAC-based or label-free quantification approaches, we assessed the impact of this difference in data composition on model performance. We therefore separated, for each modification, the data based on whether the study included SILAC-labeled spectra (wSILAC) or not (woSILAC) into two subsets as shown in table 1. Each subset was then split into training/validation/test sets as described above, using the same 85/5/10 ratio for acetylation and 89/1/10 for ubiquitination. For each subset, AHLF was fine-tuned, resulting in four additional models: AHLFa_wSILAC,

AHLFa_woSILAC, AHLFu_wSILAC and AHLFu_woSILAC. Each model was evaluated on its own test set as well as on the respective other labeled test set, e.g. the model trained on woSILAC was tested on both the woSILAC and wSILAC test sets. Additionally, a combined test dataset containing both SILAC and label-free spectra was sampled to an equal size of SILAC and label-free spectra to enable direct comparison across models.

### Code and data availability

To perform the described steps we ran custom Python (version 3.8.18) scripts, using mainly the packages NumPy (version: 1.23.5), Pandas (Version: 1.4.4), Tensorflow (Version: 2.4.1) and Pyteomics (Version: 4.7.5). The Code ran on a Linux Cluster on a single Tesla V100-SXM2-32GB GPU. All code and fine-tuned models are publicly available under our GitLab Repository(https://gitlab.com/dacs-hpi/ahlf-ptmai). The raw mass spectrometry data and search results have been deposited to the ProteomeXchange Consortium via the PRIDE (https://www.ebi.ac.uk/pride) [45] partner repository with the dataset identifier PXD079218.

## Results and Discussion

We present a curated AI ready acetylation dataset and two binary PTM classifiers for acetylation and ubiquitination based on transfer-learning from a phosphorylation-trained AHLF model. We further present an assessment of the data requirements and diversity needed for effective transfer learning for future less-studied PTMs.

### Data Harmonisation: curation and processing of human acetylation datasets

The following tables summarize the PSM counts for each dataset, along with 1% FDR, Acetyl-peptide counts, site counts, and FLR estimates. Acetylated peptidoform-site counts spanned from 2,308 (PXD023084) to 70,648 (PXD005903), with detailed metrics in Table 2.

**Table 2:** PSM Counts for Each Data Set at 1% FDR, Acetyl-peptide PSM Counts at 1% FDR, Site Counts (Excluding pA Decoy Sites) for all PSMs Collapsed by Peptidoform Position and at Each of the FLR Thresholds; 1% and 5%.

| Original data set identifier | 1% FDR PSM count | 1% FDR Acetyl-peptide count | Peptidoform-site count | 1% FLR Peptidoform site count | 5% FLR Peptidoform site count |
| --- | --- | --- | --- | --- | --- |
| PXD003530 | 274,764 | 45,256 | 14,714 | 10,901 | 12,594 |
| PXD005252 | 182,488 | 41,505 | 17,266 | 11,372 | 14,844 |
| PXD005903 | 1,768,180 | 978,629 | 70,648 | 24,116 | 49,928 |
| PXD009839 | 801,980 | 48,238 | 7,741 | 5,712 | 7,709 |
| PXD009994 | 681,833 | 230,217 | 51,762 | 36,313 | 46,536 |
| PXD023084 | 108,297 | 47,385 | 2,308 | 1,934 | 2,308 |
| PXD023218 | 359,251 | 54,610 | 6,494 | 2,134 | 4,664 |
| PXD041171 | 230,001 | 37,297 | 7,813 | 5,092 | 7,813 |
| PXD014453 | 35,774 | 17,356 | 6,926 | 2,993 | 6,926 |

Table 3 summarizes protein site counts for acetylation datasets, categorized into Gold, Silver, and Bronze tiers for lysine (K) and decoy target Alanine (A) residues. For the High-confidence (Gold) sites: Acetylated datasets yielded 8,399 Gold K sites; and Gold K-to-Bronze K ratio is 1:3.5. The predominance of Bronze K sites (29,631) highlights challenges in achieving consistent localization confidence, likely due to variable experimental conditions (see Table 1, chemical acetylation).

**Table 3:** Count for Protein Sites for Acetyl datasets, Per Category for Gold, Silver, and Bronze. G2S1B Protein Site counts in different categories (mapped to single protein)

|  | Gold K | Silver K | Bronze K | Gold A | Silver A | Bronze A |
| --- | --- | --- | --- | --- | --- | --- |
| Acetyl Builds | 8,399 | 17,555 | 29,631 | 11 | 209 | 1,886 |

### Filter datasets to apply transfer learning

After processing and filtering the acetylation datasets, the nine projects yielded a total of 840,865 modified PSMs covering 67,629 modified peptides mapping to 7,629 protein IDs. Out of those, 552,650 PSMs were either Gold or Silver labeled and therefore used for fine-tuning the acetylation model. The ubiquitination dataset contained, after processing and filtering, a total of 2,408,762 modified PSMs. Those PSMs cover 122,328 modified peptides mapping to 10,730 protein IDs. Out of those PSMs 2,277,048 are either Gold or Silver and therefore used for transfer learning. An overview and total numbers for the acetylation and ubiquitination datasets can be found in Suppl. Table S1.

### Training and Model performance

To provide PTM classification models for acetylation and ubiquitination we fine-tuned the pre-trained AHLF phosphorylation model. With heavily unbalanced datasets, 17.8% non-acetyl of all PSMs and 23.5% non-ubiquitin of all PSMs, we applied sampling strategies within our splitting to ensure data balancing and prevent data leakage. First, 183,770 sequence-overlapping contigs of modified and unmodified PSMs were first split into train, test, and validation sets. The modified PSMs thereby played the determining factor and unmodified PSMs were then sampled from within respective contigs and contigs without modified peptides to balance each set. Overall, this provided roughly 4 million training, 550,000 test, and 45,000 validation PSMs with all ubiquitinations included. The ubiquitin fine-tuned AHLF model (AHLFu) achieved a balanced accuracy (Bacc) of 0.82, and F1-score of 0.82, and an AUC of 0.90 (see Table 4). Compared to the published phosphorylation AHLFp model (median Bacc 0.77, median F1-score 0.83, median AUC 0.88) the fine-tuned ubiquitination model performs generally slightly better. This may be because of the large mass shift of the ubiquitination introduced in the spectrum. On the other hand, 115,471 contigs from acetylation resulted in a split of around 0.9 million training, 110,000 test, and 45,000 validation PSMs with all acetylations included. The Bacc of 0.78, F1-score of 0.76, and AUC of 0.87 (see Table 4) of the acetylation AHLF model (AHLFa) highlight that the performance is just slightly below the phosphorylation model. However, while acetylation results in a smaller mass shift in the spectrum than phosphorylation, a likely explanation for the deviation is the vast difference in dataset size at less than one quarter of the ubiquitination training set.

**Table 4:** Performance of the fine-tuned ubiquitination and acetylation models. The single unmod samples models, which included all modified peptides from the split configs show consistent performance with the pretrained alpha AHLFp model. The 5 models trained on full resampled training sets show overall consistent performance with the unmod samples models and additionally result in low score variance across the 5 ubiquitin and acetylation models.

| model | sampling | Bacc | F1 | AUC |
| --- | --- | --- | --- | --- |
| AHLFu | unmod | 0.82 | 0.82 | 0.90 |
| AHLFa | unmod | 0.78 | 0.76 | 0.87 |
| AHLFu | full | $0.81 \pm 0.00$ | $0.81 \pm 0.00$ | $0.89 \pm 0.00$ |
| AHLFa | full | $0.76 \pm 0.01$ | $0.74 \pm 0.02$ | $0.85 \pm 0.01$ |

### Data-saturation in Transfer-learning models

To further test the influence of the dataset size on transfer models, we adapted our sampling approach to only include the same amount of modified and unmodified PSMs from the same contig. At the same time, we continuously halved the dataset size ranging from the full set to a fraction of 1/512. Additionally, to evaluate model independence from the sampling, we performed each sampling 5 times resulting in a full set (training, test, and validation) of 478,070 *±* 753 PSMs for acetylation and 2,316,740 *±* 20 PSMs for ubiquitination, respectively with smaller sample set sizes containing as few as 935.6 *±* 0.9 and 4,526.4 *±*1847.9 PSMs, respectively. A full listing of all subsamples and splits is provided in Supplementary Table S2. Interestingly, while the entire full sampled set is roughly 50% of the entire unmod sampled dataset, the AHLFu achieved a mean Bacc of 0.81 *±* 0.00, mean F1-score of 0.81 *±* 0.00, and a mean AUC of 0.89 *±* 0.00 on the ubiquitin set, performing similarly well as the initial entire-set AHLFu model. The AHLFa model for the full sampled acetylations, on the other hand, resulted in a mean Bacc of 0.76 *±* 0.01, a mean F1-score of 0.74 *±* 0.02, and a mean AUC of 0.85 *±* 0.01, performing slightly worse than the entire-set AHLFa model (see Table 4). This highlights that even at a reduction of the dataset the transfer-learning approach is successful and a valuable tool for further modification assessments. Furthermore, the low standard deviations across both models indicate that the performance does not depend on the individual sampling splits, asserting stable performance of the training and models.

To evaluate how much training data is required to effectively perform transfer-learning, we trained additional models for the further down-sampled sets and compared their results (see Figure 2 and Figure 3).

**Figure 2:**
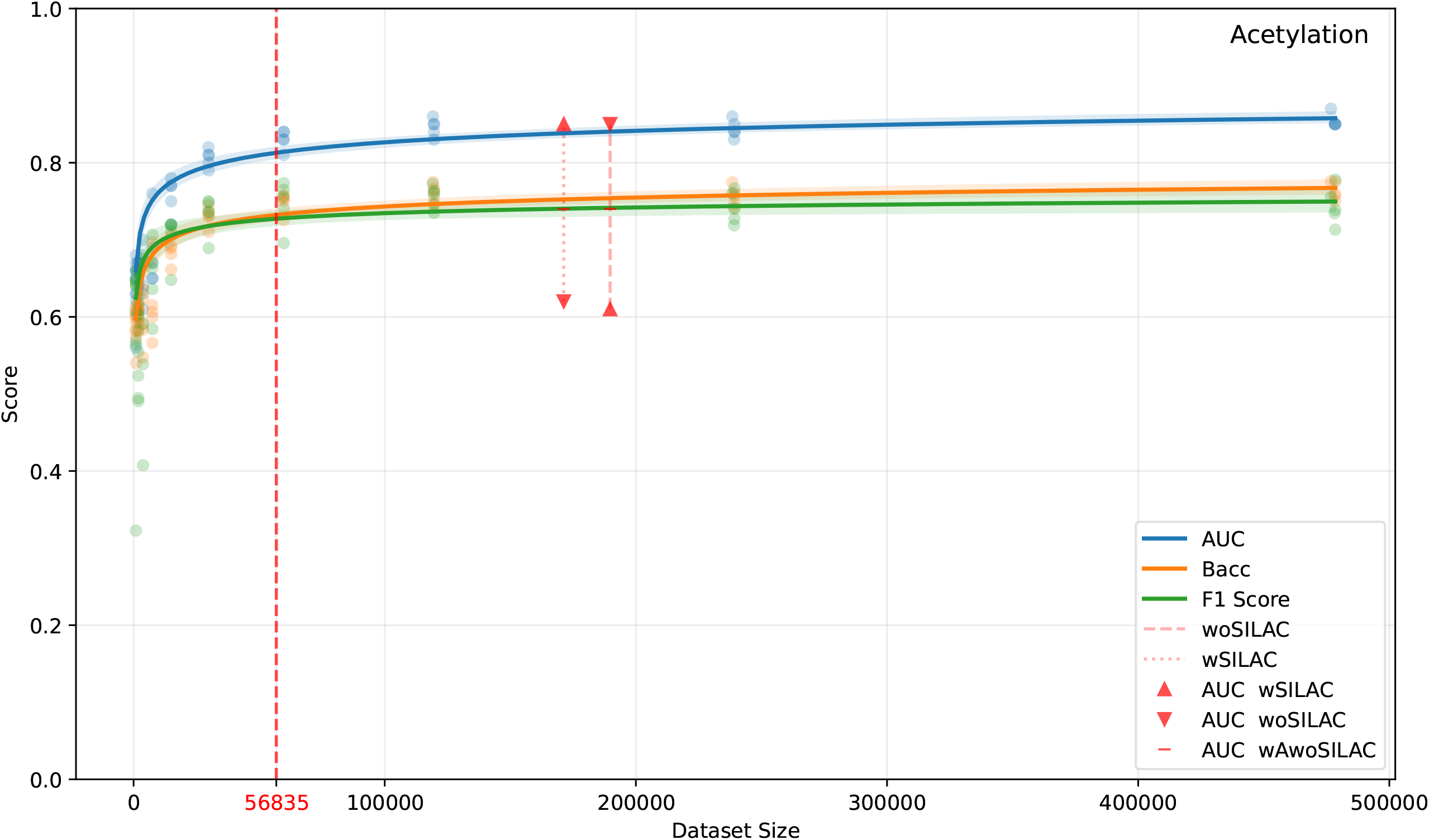
Data scaling curves for the acetylation model showing Bacc, F1 and AUC across iteratively down-sampled training set sizes, fitted using a power law function. The red dashed vertical line marks the elbow point at 57,000 PSMs, below which model performance drops substantially. Horizontal dotted lines indicate the AUC of models trained on pure SILAC (wSILAC) and pure label-free (woSILAC) data, with triangles marking their respective training dataset sizes and short dashes indicating performance on the combined test set (wA-woSILAC)

**Figure 3:**
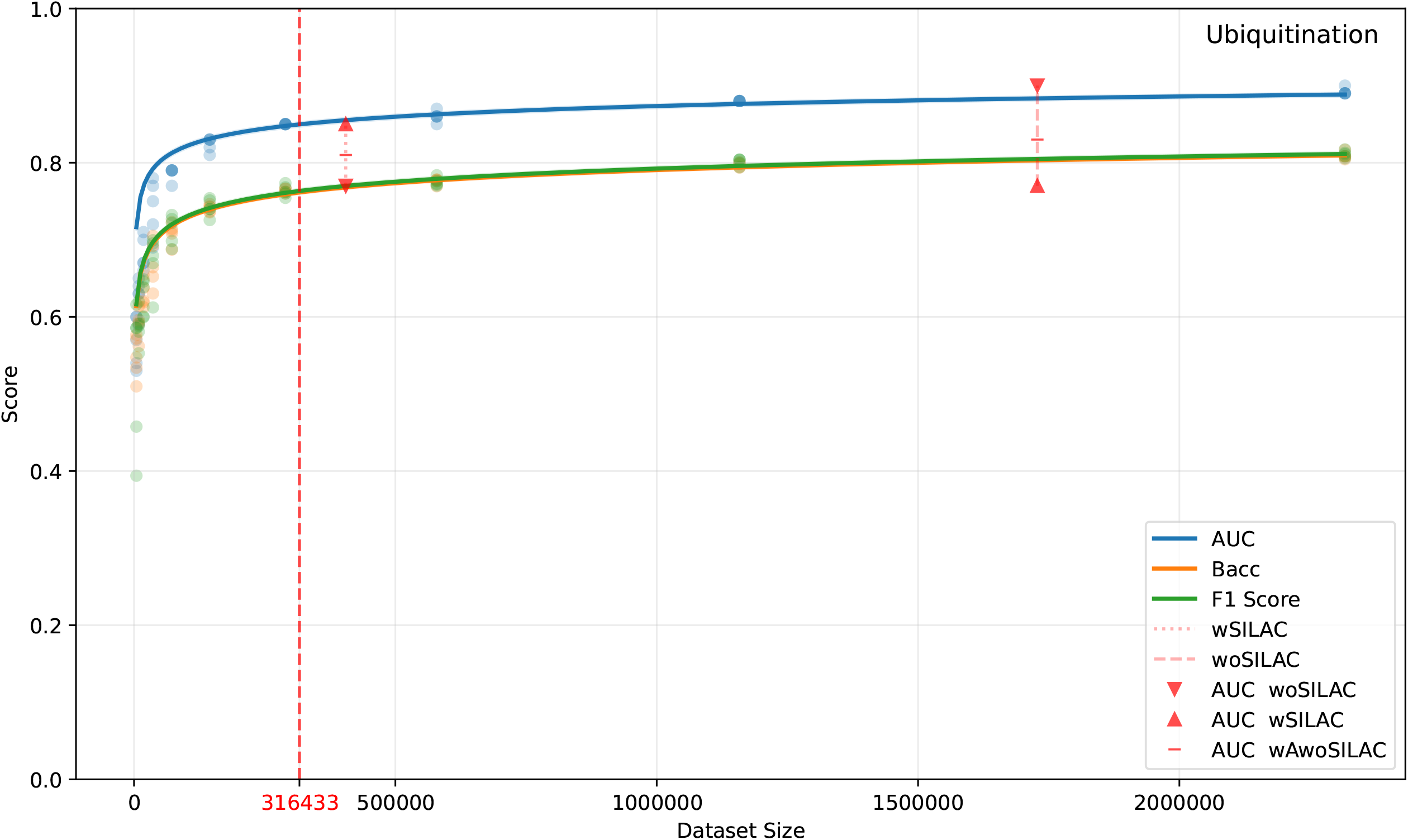
Data scaling curves for the ubiquitination model showing Bacc, F1 and AUC across iteratively down-sampled training set sizes, fitted using a power law function. The red dashed vertical line marks the elbow point at 316,433 PSMs, below which model performance drops substantially. Horizontal dotted lines indicate the AUC of models trained on pure SILAC (wSILAC) and pure label-free (woSILAC) data, with triangles marking their respective training dataset sizes and short dashes indicating performance on the combined test set (wAwoSILAC).

For acetylation we were able to fit a power law function for the different metrics across the dataset sizes, as illustrated in Figure 2. The performance degrades only gradually with initial reductions in dataset size, until the elbow point at approximately 57,000 PSMs, corresponding to 28,500 acetylated PSMs. Below this point the performance of the trained model drops substantially, marking the minimum dataset size recommended for the acetylation model.

Below this 1/8th (12.5%) point, having fewer PSMs drastically impacts the model performance to the point of the classifier merely outperforming random guessing. Similarly, the ubiquitination models follow the same trajectory and highlight an elbow point at a down-sampling of roughly 1/8th (12.5%) as shown in Figure 3. However, due to the initially larger dataset size, ubiquitination requires roughly 316,000 PSMs (158,000 PSMs with a ubiquitination) before additional data provide diminishing improvements.

The substantially higher data requirement compared to acetylation suggests that the model requires more data to learn discriminative features for ubiquitination, which could reflect the nature of the modification itself, which is only detectable through the two additional glycines. Overall, for both modifications the flattening curve in each metric for subsamples closer in size to the original sets highlight that our original datasets are well within an optimal size for the transfer-learning as not much could be gained with larger datasets. This is particularly relevant given the scarcity of high quality PTM data [9]. Especially, the relatively modest data requirement for acetylation encourages the application for other PTMs where data availability is even more limited, as was shown for methylation [46]. The findings of this work provide a practical guideline for researchers to extend PTM classification to new modification types using transfer learning.

### Impact of data diversity

All projects used for the curation of both datasets were selected and processed similarly even though the quantification method differs within each modification subset (see Supplementary Table 1). Label-free and SILAC labelling, specifically the light versions do not significantly differ in their MS/MS representation. However, minor differences may occur that so far have not been observed. Therefore, we trained four additional AHLF models to investigate the impact of data diversity. The data were separated into SILAC and label-free datasets for both acetylation and ubiquitination and split into training, test and validation sets. Separate models were fine-tuned on each pure training subset. As expected, models performed best, when they were evaluated on their respective test sets. AHFLa_wSILAC achieved a Bacc of 0.77 and an AUC of 0.85 on the SILAC test set, while AHLFa_woSILAC achieved a Bacc of 0.77 and an AUC of 0.85 on the label-free test set (see Figure 2). A similar pattern can be observed for ubiquitination. AHLFu_wSILAC achieves a Bacc of 0.77 and an AUC of 0.85 on SILAC data and AHLFu_woSILAC achieving a Bacc of 0.81 and an AUC of 0.90 on label free data highlighted in figure 3. Additionally, the results are within the 95% confidence intervals for similar sized mixed-quantification type models. This is consistent with the expectation that more uniform training data reduces the complexity the model needs to handle, by removing one layer of diversity from the data which the model needs to handle. We then created two combined test sets, one for each PTM, to contain balanced amounts of SILAC and label-free PSMs and assessed the respective PTM model with them (see Figure 2, Figure 3, and Table 2).

For acetylation, both AHLFa_wSILAC and AHLFa_woSILAC achieved an AUC of 0.74 on the combined set. For ubiquitination, AHLFu_wSILAC achieved an AUC of 0.81 and AHLFu_woSILAC achieved an AUC of 0.83. As can be observed in Figure 2 and Figure 3 the models tested on their respective matching test set, that is the models trained wSILAC and tested wSILAC and models trained woSILAC and tested woSILAC, with exception of the AHLFu_wSILAC model, sit above the scaling AUC curves at their respective dataset size. This suggests that training on clean uniform data allows to slightly outperform an equivalent mixed model on mixed data. However, the performance drop on the combined test set reveals that pure models do not generalize well across data types. When evaluating the models on the opposite test data the performance dropped even more substantially (see Figures 2, Figure 3, and Table 5). For acetylation AHLFa_wSILAC dropped to an AUC of 0.62 on label-free data (woSILAC) and AHLFa_woSILAC dropped to an AUC of 0.61 on SILAC data (wSILAC). Also for ubiquitination the performance dropped notably. So AHLFu_wSILAC dropped to an AUC of 0.77 on the label-free data (woSILAC) and AHLFu_woSILAC dropped to an AUC of 0.77 on SILAC data (wSILAC). This indicates that the models tend to overfit to the spectral characteristics of the respective training data type. Overall, the results highlight that a mixed dataset between label-free and SILAC data combined draws not only from the benefit of a larger dataset, but also from a better combined representation of potential small differences introduced by the modification and even more substantial differences by the heavy SILAC labeled sequences., This highlights that the transfer-learning approach based on the originally learned spectrum representation in the AHLFp model accommodates modification as targets and at the same time learns differences due to labelling as part of the representation.

**Table 5:** Evaluation of the impact of data diversity in the model performance. Rows represent the models trained only with SILAC and without SILAC data for acetylation and ubiquitination, respectively. Columns highlight the different metrics based on the applied three test data cases: with SILAC, without SILAC, and all combined.

|  | with SILAC (wSILAC) |  |  | label free (woSILAC) |  |  | combined (wAwoSILAC) |  |  |
| --- | --- | --- | --- | --- | --- | --- | --- | --- | --- |
|  | Bacc | F1 | AUC | Bacc | F1 | AUC | Bacc | F1 | AUC |
| AHLFa_wSILAC | 0.77 | 0.76 | 0.85 | 0.58 | 0.50 | 0.62 | 0.68 | 0.64 | 0.74 |
| AHLFa_woSILAC | 0.57 | 0.58 | 0.61 | 0.77 | 0.77 | 0.85 | 0.67 | 0.67 | 0.74 |
| AHLFu_wSILAC | 0.77 | 0.77 | 0.85 | 0.69 | 0.73 | 0.77 | 0.73 | 0.74 | 0.81 |
| AHLFu_woSILAC | 0.69 | 0.71 | 0.77 | 0.81 | 0.81 | 0.90 | 0.75 | 0.76 | 0.83 |

## Conclusion

Curation and harmonisation of AI ready datasets from public repositories provides opportunities but also suffers from challenges. Our harmonisation of an human acetylation dataset highlights the challenges from identifying suitable public datasets, consistent reanalysis and curation of identifications. At the same time, we found that datasets large enough for machine-learning tasks such as transfer-learning can be achieved for PTMs that are less studied compared to phosphorylation. We showed that applying transfer learning on a model capable of classifying phosphorylated spectra is an effective strategy to extend PTM classification to new modification types. To this end, we successfully developed two new models for acetylation and ubiquitination. In addition we showed by reducing the amount of training data, that fine-tuning requires a modest amount of data depending on the changes to the spectrum introduced by the modification, with around 28,500 modified spectra for acetylation and 158,000 for ubiquitination. This is encouraging for future extensions to other PTMs, with fewer public enrichment datasets available for curation. Furthermore, we highlight that training on diverse data, by combining SILAC and label-free spectra, is a beneficial strategy for generalizability, while pure models tend to overfit to their data type. The framework presented in this study could be extended to other PTM types where sufficient data is available or can be curated following the approach presented here.

## Supplemental Data

Supplementary Table S1: Overview of datasets including number of peptide-spectrum matches, modifications and FLR category.

Supplementary Table S2: Overview of model performances for down-sampled acetylation and ubiqituitination subsets including dataset fraction, size, and model performance metrics.

## References

(1) Pan, S.; Chen, R. Pathological implication of protein post-translational modifications in cancer, Mol. Asp. Med. (2022), 86, 101097, DOI: 10.1016/j.mam.2022.101097.

(2) Wu, X.; Xu, M.; Geng, M.; Chen, S.; Little, P. J.; Xu, S.; Weng, J. Targeting protein modifications in metabolic diseases: molecular mechanisms and targeted therapies, Signal Transduct. Target. Ther. (2023), 8, 220, DOI: 10.1038/s41392-023-01439-y.

(3) Kumar, M.; Schlaffner, C. N.; Tang, S.; Beuvink, M. A.; Viode, A.; Mair, W.; Jha, M.; Uncu, C.; Wesseling, H.; Wang, T.; Oakley, D. H.; Beerepoot, P.; Xue, J.; Connors, T. R.; Davis, D. A.; Frosch, M. P.; Murray, M. E.; Spina, S. E.; Grinberg, L. T.; Seeley, W. W.; Miller, B. L.; Boxer, L.; Geschwind, D. H.; Kosik, K. S.; Dickson, D. W.; Renard, B. Y.; DeTure, M.; McKee, A. C.; Hyman, B. T.; Steen, H.; Steen, J. A. Molecular features of human pathological tau distinguish tauopathy-associated dementias, Cell (2026), 189, 956–968.e13, DOI: 10.1016/j.cell.2025.12.036.

(4) Zhong, Q.; Xiao, X.; Qiu, Y.; Xu, Z.; Chen, C.; Chong, B.; Zhao, X.; Hai, S.; Li, S.; An, Z.; Dai, L. Protein posttranslational modifications in health and diseases: Functions, regulatory mechanisms, and therapeutic implications, MedComm (2023), 4, e261, DOI: 10.1002/mco2.261.

(5) Kitamura, N.; Galligan, J. J. A global view of the human post-translational modification landscape, Biochem. J. (2023), 480, 1241–1265, DOI: 10.1042/BCJ20220251.

(6) Wu, F.; Li, W.; Lu, H.; Li, L. Recent Advances in Mass Spectrometry-Based Studies of Post-Translational Modifications in Alzheimer’s Disease, Mol. Cell. Proteom. (2025), 24, 101003, DOI: 10.1016/j.mcpro.2025.101003.

(7) Lai, S.; Dai, S.; Zhao, P.; Zhou, C.; Li, N.; Yu, W. PIPI-C: A Combinatorial Optimization Framework for Identifying Post-translational Modification Hot-spots in Mass Spectrometry Data, Mol. Cell. Proteom. (2026), 25, 101494, DOI: 10.1016/j.mcpro.2025.101494.

(8) Kertész-Farkas, A.; Reiz, B.; Vera, R.; Myers, M. P.; Pongor, S. PTMTreeSearch: a novel two-stage tree-search algorithm with pruning rules for the identification of post-translational modification of proteins in MS/MS spectra, Bioinformatics (2014), 30, 234–241, DOI: 10.1093/bioinformatics/btt642.

(9) Bittremieux, W.; Ananth, V.; Fondrie, W. E.; Melendez, C.; Pominova, M.; Sanders, J.; Wen, B.; Yilmaz, M.; Noble, W. S. Deep Learning Methods for De Novo Peptide Sequencing, Mass Spectrom. Rev. (2024), DOI: 10.1002/mas.21919.

(10) Straub, G.; Ananth, V.; Fondrie, W. E.; Hsu, C.; Klaproth-Andrade, D.; Pominova, M.; Riffle, M.; Sanders, J.; Wen, B.; Xu, L.; Yilmaz, M.; MacCoss, M. J.; Oh, S.; Bittremieux, W.; Noble, W. S. Improvements to Casanovo, a Deep Learning De Novo Peptide Sequencer, J. Proteome Res. (2026), 25, 755–764, DOI: 10.1021/acs.jproteome.5c00706.

(11) Klaproth-Andrade, D.; Bruns, Y.; Gabriel, W.; Nix, C.; Bergant, V.; Pichlmair, A.; Wilhelm, M.; Gagneur J. Modanovo: A Unified Model for Post-translational Modification-Aware De Novo Sequencing Using Experimental Spectra From In Vivo and Synthetic Peptides, Mol. Cell. Proteom. (2026), 25, 101501, DOI: 10.1016/j.mcpro.2025.101501.

(12) Yilmaz, M.; Fondrie, W.; Bittremieux, W.; Oh, S.; Noble, W. S. In Proceedings of the 39th International Conference on Machine Learning, ed. by Chaudhuri, K.; Jegelka, S.; Song, L.; Szepesvari, C.; Niu, G.; Sabato, S., PMLR: 2022; Vol. 162, pp 25514–25522.

(13) Altenburg, T.; Giese, S. H.; Wang, S.; Muth, T.; Renard, B. Y. Ad hoc learning of peptide fragmentation from mass spectra enables an interpretable detection of phosphorylated and cross-linked peptides, Nat. Mach. Intell. (2022), 4, 378–388, DOI: 10.1038/s42256-022-00467-7.

(14) Dens, C.; Adams, C.; Laukens, K.; Bittremieux, W. Machine Learning Strategies to Tackle Data Challenges in Mass Spectrometry-Based Proteomics, J. Am. Soc. Mass Spectrom. (2024), 35, 2143–2155, DOI: 10.1021/jasms.4c00180.

(15) Wang, M.; Wang, J.; Carver, J.; Pullman, B. S.; Cha, S. W.; Bandeira, N. Assembling the Community-Scale Discoverable Human Proteome, Cell Syst. (2018), 7, 412–421.e5, DOI: 10.1016/j.cels.2018.08.004.

(16) Gabriel, W.; Shouman, O.; Schroeder, A.; Boessl, F.; Wilhelm, M. In Advances in Neural Information Processing Systems, ed. by Globerson, A.; Mackey, L.; Belgrave, D.; Fan, A.; Paquet, U.; Tomczak, J.; Zhang, C., Curran Associates, Inc.: 2024; Vol. 37, pp 131154–131196, DOI: 10.52202/079017-4168.

(17) Zolg, D. P.; Wilhelm, M.; Schmidt, T.; Médard, G.; Zerweck, J.; Knaute, T.; Wenschuh, H.; Reimer, U.; Schnatbaum, K.; Kuster, B. ProteomeTools: Systematic Characterization of 21 Post-translational Protein Modifications by Liquid Chromatography Tandem Mass Spectrometry (LC-MS/MS) Using Synthetic Peptides*, Mol. Cell. Proteom. (2018), 17, 1850–1863, DOI: 10.1074/mcp.TIR118.000783.

(18) An, Z.; Zhai, L.; Ying, W.; Qian, X.; Gong, F.; Tan, M.; Fu, Y. PTMiner: Localization and Quality Control of Protein Modifications Detected in an Open Search and Its Application to Comprehensive Post-translational Modification Characterization in Human Proteome, Mol. Cell. Proteom. (2019), 18, 391–405, DOI: 10.1074/mcp.RA118.000812.

(19) Deutsch, E. W.; Bandeira, N.; Perez-Riverol, Y.; Sharma, V.; Carver, J. J.; Mendoza, L.; Kundu, D. J.; Bandla, C.; Kamatchinathan, S.; Hewapathirana, S.; Sun, Z.; Kawano, S.; Okuda, S.; Connolly, B.; MacLean, B.; MacCoss, M. J.; Chen, T.; Zhu, Y.; Ishihama, Y.; Vizcaíno, J. A. The ProteomeXchange consortium in 2026: making proteomics data FAIR, Nucleic Acids Res. (D1 2026), 54, D459–D469, DOI: 10.1093/nar/gkaf1146.

(20) Ochoa, D.; Jarnuczak, A. F.; Viéitez, C.; Gehre, M.; Soucheray, M.; Mateus, A.; Kleefeldt, A.; Hill, A.; Garcia-Alonso, L.; Stein, F.; Krogan, N. J.; Savitski, M. M.; Swaney, D. L.; Vizcaíno, J. A.; Noh, K.-M.; Beltrao, P. The functional landscape of the human phosphoproteome, Nat. Biotechnol. (2020), 38, 365–373, DOI: 10.1038/s41587-019-0344-3.

(21) Ramazi, S.; Zahiri, J. Post-translational modifications in proteins: resources, tools and prediction methods, Database (2021), 2021, baab012, DOI: 10.1093/database/baab012.

(22) Li, Y.-Y.; Liu, Z.; Liu, X.; Zhu, Y.-H.; Fang, C.; Arif, M.; Qiu, W.-R. A Systematic Review of Computational Methods for Protein Post-Translational Modification Site Prediction, Arch. Comput. Methods Eng. (2025), DOI: 10.1007/s11831-025-10444-z.

(23) Yau, R.; Rape, M. The increasing complexity of the ubiquitin code, Nat. Cell Biol. (2016), 18, 579–586, DOI: 10.1038/ncb3358.

(24) Wang, W. W.; Singha Roy, S. J.; Parker, C. G. Targeted Protein Acetylation Through Chemically Induced Proximity, Acc. Chem. Res. (2025), 58, 2695–2707, DOI: 10.1021/acs.accounts.5c00326.

(25) Gerwen, J. v.; Fottner, M.; Wang, S.; Busby, B.; Boswell, E.; Schnacke, P.; Carrano, A. C.; Bakowski, M. A.; Troemel, E. R.; Studer, R.; Strumillo, M.; Martin, M.-J.; Harper, J. W.; Lang, K.; Jones, A. R.; Bennett, E. J.; Vizcaíno, J. A.; Barrio-Hernandez, I.; Beltrao, P. Mapping the functional importance of site-specific ubiquitination across the human proteome, 2026, DOI: 10.1101/2025.10.08.681129.

(26) Tatham, M. H.; Cole, C.; Scullion, P.; Wilkie, R.; Westwood, N. J.; Stark, L. A.; Hay, R. T. A Proteomic Approach to Analyze the Aspirin-mediated Lysine Acetylome*, Mol. Cell. Proteom. (2017), 16, 310–326, DOI: 10.1074/mcp.O116.065219.

(27) Weinert, B. T.; Narita, T.; Satpathy, S.; Srinivasan, B.; Hansen, B. K.; Schölz, C.; Hamilton, W. B.; Zucconi, B. E.; Wang, W. W.; Liu, W. R.; Brickman, J. M.; Kesicki, E. A.; Lai, A.; Bromberg, K. D.; Cole, P. A.; Choudhary, C. Time-Resolved Analysis Reveals Rapid Dynamics and Broad Scope of the CBP/p300 Acetylome, Cell (2018), 174, 231–244.e12, DOI: 10.1016/j.cell.2018.04.033.

(28) Gil, J.; Ramírez-Torres, A.; Chiappe, D.; Luna-Peñaloza, J.; Fernandez-Reyes, F. C.; Arcos-Encarnación, B.; Contreras, S.; Encarnación-Guevara, S. Lysine acetylation stoichiometry and proteomics analyses reveal pathways regulated by sirtuin 1 in human cells, J. Biol. Chem. (2017), 292, 18129–18144, DOI: 10.1074/jbc.M117.784546.

(29) Murray, L. A.; Sheng, X.; Cristea, I. M. Orchestration of protein acetylation as a toggle for cellular defense and virus replication, Nat. Commun. (2018), 9, 4967, DOI: 10.1038/s41467-018-07179-w.

(30) Hansen, B. K.; Gupta, R.; Baldus, L.; Lyon, D.; Narita, T.; Lammers, M.; Choudhary, C.; Weinert, B. T. Analysis of human acetylation stoichiometry defines mechanistic constraints on protein regulation, Nat. Commun. (2019), 10, 1055, DOI: 10.1038/s41467-019-09024-0.

(31) Hostrup, M.; Lemminger, A. K.; Stocks, B.; Gonzalez-Franquesa, A.; Larsen, J. K.; Quesada, J. P.; Thomassen, M.; Weinert, B. T.; Bangsbo, J.; Deshmukh, A. S. High-intensity interval training remodels the proteome and acetylome of human skeletal muscle, eLife (2022), 11, ed. by James, D. E.; Kuroda, S., e69802, DOI: 10.7554/eLife.69802.

(32) Zecha, J.; Gabriel, W.; Spallek, R.; Chang, Y.-C.; Mergner, J.; Wilhelm, M.; Bassermann, F.; Kuster, B. Linking post-translational modifications and protein turnover by site-resolved protein turnover profiling, Nat. Commun. (2022), 13, 165, DOI: 10.1038/s41467-021-27639-0.

(33) Betsinger, C. N.; Justice, J. L.; Tyl, M. D.; Edgar, J. E.; Budayeva, H. G.; Abu, Y. F.; Cristea, I. M. Sirtuin 2 promotes human cytomegalovirus replication by regulating cell cycle progression, mSystems (2023), 8, e00510–23, DOI: 10.1128/msystems.00510-23.

(34) Baeza, J.; Lawton, A. J.; Fan, J.; Smallegan, M. J.; Lienert, I.; Gandhi, T.; Bernhardt, O. M.; Reiter, L.; Denu, J. M. Revealing Dynamic Protein Acetylation across Subcellular Compartments, J. Proteome Res. (2020), 19, 2404–2418, DOI: 10.1021/acs.jproteome.0c00088.

(35) Hulstaert, N.; Shofstahl, J.; Sachsenberg, T.; Walzer, M.; Barsnes, H.; Martens, L.; Perez-Riverol, Y. ThermoRawFileParser: Modular, Scalable, and Cross-Platform RAW File Conversion, J. Proteome Res. (2020), 19, 537–542, DOI: 10.1021/acs.jproteome.9b00328.

(36) Eng, J. K.; Jahan, T. A.; Hoopmann, M. R. Comet: An open-source MS/MS sequence database search tool, Proteomics (2013), 13, 22–24, DOI: 10.1002/pmic.201200439.

(37) Keller, A.; Nesvizhskii, A. I.; Kolker, E.; Aebersold, R. Empirical Statistical Model To Estimate the Accuracy of Peptide Identifications Made by MS/MS and Database Search, Anal. Chem. (2002), 74, 5383–5392, DOI: 10.1021/ac025747h.

(38) Shteynberg, D.; Deutsch, E. W.; Lam, H.; Eng, J. K.; Sun, Z.; Tasman, N.; Mendoza, L.; Moritz, R. L.; Aebersold, R.; Nesvizhskii, A. I. iProphet: Multi-level Integrative Analysis of Shotgun Proteomic Data Improves Peptide and Protein Identification Rates and Error Estimates*, Mol. Cell. Proteom. (2011), 10, M111.007690, DOI: 10.1074/mcp.M111.007690.

(39) Deutsch, E. W.; Mendoza, L.; Shteynberg, D. D.; Hoopmann, M. R.; Sun, Z.; Eng, J. K.; Moritz, R. L. Trans-Proteomic Pipeline: Robust Mass Spectrometry-Based Proteomics Data Analysis Suite, J. Proteome Res. (2023), 22, 615–624, DOI: 10.1021/acs.jproteome.2c00624.

(40) Shteynberg, D. D.; Deutsch, E. W.; Campbell, D. S.; Hoopmann, M. R.; Kusebauch, U.; Lee, D.; Mendoza, L.; Midha, M. K.; Sun, Z.; Whetton, A. D.; Moritz, R. L. PTMProphet: Fast and Accurate Mass Modification Localization for the Trans-Proteomic Pipeline, J. Proteome Res. (2019), 18, 4262–4272, DOI: 10.1021/acs.jproteome.9b00205.

(41) Ramsbottom, K. A.; Prakash, A.; Riverol, Y. P.; Camacho, O. M.; Martin, M.-J.; Vizcaíno, J. A.; Deutsch, E. W.; Jones, A. R. Method for Independent Estimation of the False Localization Rate for Phosphoproteomics, J. Proteome Res. (2022), 21, 1603–1615, DOI: 10.1021/acs.jproteome.1c00827.

(42) Ramsbottom, K. A.; Prakash, A.; Perez-Riverol, Y.; Camacho, O. M.; Sun, Z.; Kundu, D. J.; Bowler-Barnett, E.; Martin, M.; Fan, J.; Chebotarov, D.; McNally, K. L.; Deutsch, E. W.; Vizcaíno, J. A.; Jones, A. R. Meta-Analysis of Rice Phosphoproteomics Data to Understand Variation in Cell Signaling Across the Rice Pan-Genome, J. Proteome Res. (2024), 23, 2518–2531, DOI: 10.1021/acs.jproteome.4c00187.

(43) Camacho, O. M.; Ramsbottom, K. A.; Collins, A.; Jones, A. R. Assessing Multiple Evidence Streams to Decide on Confidence for Identification of Post-Translational Modifications, within and Across Data Sets, J. Proteome Res. (2023), 22, 1828–1842, DOI: 10.1021/acs.jproteome.2c00823.

(44) Satopa, V.; Albrecht, J.; Irwin, D.; Raghavan, B. In 31st International Conference on Distributed Computing Systems Workshops, 2011, pp 166–171.

(45) Perez-Riverol, Y.; Bai, J.; Bandla, C.; García-Seisdedos, D.; Hewapathirana, S.; Kamatchinathan, S.; Kundu, D. J.; Prakash, A.; Frericks-Zipper, A.; Eisenacher, M.; Walzer, M.; Wang, S.; Brazma, A.; Vizcaíno, J. A. The PRIDE database resources in 2022: a hub for mass spectrometry-based proteomics evidences, Nucleic Acids Res. (D1 2022), 50, D543–D552, DOI: 10.1093/nar/gkab1038.

(46) Wang, S.; Hartmaring, Y.; Schlaffner, C. N.; Bowler-Barnett, E. H.; Martin, M.; Fan, J.; Sun, Z.; Renard, B. Y.; Deutsch, E. W.; Jones, A. R.; Vizcaíno, J. A. A High-Confidence Atlas of Protein Methylation Enables AI-Driven Detection of Methylated Peptides, bioRxiv (2026), DOI: 10.64898/2026.07.01.733993.

